# Crosstalk of the MAP3K1 and EGFR pathways mediates gene-environment interactions that disrupt developmental tissue closure

**DOI:** 10.1101/2024.03.14.585101

**Authors:** Jingjing Wang, Bo Xiao, Eiki Kimura, Maureen Mongan, Wei-Wen Hsu, Mario Medvedovic, Alvaro Puga, Ying Xia

## Abstract

Aberrant signal transduction pathways can adversely derail developmental processes. One such process is embryonic eyelid closure that requires MAP3K1. *Map3k1* knockout mice have defective eyelid closure and an autosomal recessive eye-open at birth phenotype. *In utero* exposure to dioxin, a persistent environmental toxicant, causes the same eye defect in *Map3k1*^+/-^ hemizygous but not wild type pups. Here we explore the mechanisms of *Map3k1* (gene) and dioxin (environment) interactions (GxE) in the tissue closure defect. We show that, acting through the AHR, dioxin activates EGFR signaling, which in turn depresses MAP3K1-dependent JNK activity. This effect of dioxin is exacerbated by *Map3k1* heterozygosity. Therefore, dioxin exposed *Map3k1^+/-^* embryonic eyelids have a marked reduction of JNK activity, accelerated differentiation and impeded polarization in the epithelial cells. Knocking out *Ahr* or *Egfr* in eyelid epithelium attenuates the open-eye defects in dioxin-treated *Map3k1^+/-^* pups, whereas knockout of *Jnk1* and *S1pr*, encoding the S1P receptors upstream of the MAP3K1-JNK pathway, potentiates dioxin toxicity. Our novel findings suggest that dioxin and genes of the AHR, EGFR and S1P-MAP3K1-JNK pathways constitute a multifactorial mechanism underlying tissue closure abnormalities.

**Summary statement:** The crosstalk between a global environmental pollutant and the pre-existing genetic conditions is mediated through interactive signaling pathways, resulting in anatomical tissue closure abnormalities in development.

## Introduction

The formation of a complex multicellular organism is one of the most fascinating processes in biology. These processes require effective transmission of developmental cues by signal transduction pathways to regulate cellular activities at the correct time and space. The Mitogen Activated Protein Kinase (MAPK) pathway is an evolutionarily conserved signaling mechanism that exerts global developmental control (Kuida and Boucher, 2004). The pathway consists of a three-tiered kinase module, including a MAPKK kinase (MAP3K), a MAPK kinase (MAP2K) and a MAPK. For each module, the endogenous and environmental signals activate the MAP3Ks that in turn phosphorylate the MAP2Ks to further phosphorylate and activate the MAPKs. The MAP3K is a family of 20 structurally related serine-threonine protein kinases that play a crucial role in determining specificity of MAPK activation (Hagemann and Blank, 2001). In this capacity, each MAP3K is responsive to a distinctive set of upstream signals and in turn activates the specific MAP2K-MAPK modules (Uhlik et al., 2004). Consistant with their specificity in signal transduction, the MAP3Ks exhibit highly cell type- and tissue-specific roles in development (Craig et al., 2008). The MAP3K1, for example, is dispensable for survival, but uniquely needed for embryonic eyelid closure (Wang et al., 2021a).

Mouse eyelid development starts at embryonic day 11.5 (E11.5), where the periocular epithelium derived from the surface ectoderm folds at the junction of the future conjunctiva and cornea, forming the eyelid buds. The eyelid buds elongate as embryo grows and the epithelia at the eyelid leading edge extend centripetally. By E15.5, the opposing eyelid epithelium makes contact and fuses with each other, resulting in an enclosed eyelid covering the ocular surface (Harris and Juriloff, 1986, Findlater et al., 1993). The mouse eyelids remain fused at birth, and the upper and lower eyelids separate, resulting in eyes open at postnatal day 12-14, a time point coinciding with developmental maturation of the major ocular surface tissues. Thus, mice are normally born with eyes closed; however, the newborn pups display an “eye-open” at birth (EOB) phenotype when the eyelids fail to fuse prenatally (Teramoto et al., 1988). Because the failure of eyelid closure does not impair prenatal survival and the EOB phenotype is distinct and easily detectable (Meng et al., 2014), mouse strains with eyelid closure defects are convenient research tools to elucidate the mechanisms of developmental tissue closure (Gage et al., 2008, Schaeper et al., 2007, Qu et al., 1999, Huang et al., 2009, Tao et al., 2006) Compelling genetic data have shown that *Map3k1* loss-of-function mutation is autosomal recessive for the EOB phenotype (Yujiri et al., 1999, Zhang et al., 2003, Juriloff et al., 2005, Parker et al., 2015). While the hemizygous mice have normal eyelid development, the homozygous knockouts exhibit an EOB defect with full penetrance. The same EOB defects are found in mice lacking MAP3K1 kinase domain, suggesting that its activity in MAPK signaling is essential for eyelid closure. In alignment with this suggestion, MAP3K1 is found required for optimal activation of the Jun N-terminal Kinases (JNK) in epithelial cells at the leading edge of the embryonic eyelids (Zhang et al., 2003, Zhang et al., 2005). Genetic testing lends further support to the existence of a MAP3K1-JNK pathway in eyelid morphogenesis. While neither the *Map3k1* heterozygosity (*Map3k1^+/-^*) nor the *Jnk1* knockout (*Jnk^-/-^*) and the *Jnk1^+/-^Jnk2^+/-^* double heterozygous mice exhibit the EOB phenotype, the *Map3k1^+/-^Jnk1^-/-^*and *Map3k1^+/-^Jnk1^+/-^Jnk2^+/-^* compound mutants have the defects (Takatori et al., 2008). The genetic non-allelic non-complementation strongly indicates that *Map3k1*, *Jnk1* and *Jnk2* gene products function in the same molecular pathway for embryonic eyelid closure (Yook et al., 2001).

TCDD (2,3,7,8-tetrachlorodibenzo-para-dioxin) is an organochlorinated pollutant and the prototype of dozens of ubiquitous environmental compounds, known collectively as dioxin-like chemicals (DLCs). These chemicals are bio-persistent and ubiquitous in the environment; thus, human exposure is inevitable. Exposure of DLC has been linked to irregularities of many crucial life functions, including damage to the immune system, cardiovascular disease, diabetes, hormone dysfunction and cancer (Chapin et al., 2004). The biological effects of TCDD are mediated by the Aromatic Hydrocarbon Receptor (AHR), a ligand activated basic-region helix–loop–helix-PAS transcription factor (Peters et al., 1999, Fernandez-Salguero et al., 1996). After binding with TCDD, the AHR translocates from the cytoplasm to the nucleus where it heterodimerizes with the aryl hydrocarbon receptor nuclear translocator (ARNT) and/or interacts with other transcription factors to, directly or indirectly, regulate gene expression (Hankinson, 1995, Fujii-Kuriyama and Mimura, 2005, Puga et al., 2000, Elferink, 2003). AHR-regulated genes are responsible for most of the developmental toxicities of TCDD (Hurst et al., 1998, Abbott, 1995, Abbott et al., 1998, Couture et al., 1990, Yoshioka et al., 2011, Fernandez-Salguero et al., 1996); however, there is evidence suggesting that genes outside of the *Ahr* gene battery can modify the toxicity, resulting in species and individual variations, and cell type-specific responses (Thomae et al., 2006, Keller et al., 2008). The modification mechanism is a trending topic of investigation to understand the complexity and individual diversity associated with TCDD toxicity.

When studying the developmental toxicity of TCDD in mice, we came cross an observation that its exposure *in utero* induced an open-eye defect in hemizygous *Map3k1^+/-^* pups. By contrast, exposure did not cause the defects in wild type pups and in pups carrying *Dkk2* loss-of-function mutations that target the WNT signaling pathways (Mongan et al., 2015). In the current work, we investigated the mechanisms of the specific interactions between gene (*Map3k1*) x environment (TCDD) (GxE) in tissue closure abnormalities. We found that the toxicity of TCDD was mediated through the AHR to activate the Epidermal Growth Factor Receptor (EGFR) - Extracellular signal Regulated Kinase (ERK) pathway. This in turn led to a modest inhibition of JNK activity that was insufficient to cause an EOB phenotype. *Map3k1* heterozygosity also decreased JNK activity, and it acted additively or synergistically with TCDD to further inactive JNK, resulting in the eyelid closure defects in TCDD-exposed *Map3k1^+/-^*pups. We additionally identified the Sphingosine 1-phosphate (S1P) receptors as upstream of the MAP3K1-JNK pathway and that knockout of *Jnk1*- or *S1pr2/3* also sensitized the embryonic eyelids to TCDD-induced closure defects. Results reported here show that *Map3k1* heterozygosity and TCDD converge to inhibit JNK signaling through intertwining crosstalk of multiple signaling pathways.

## Results

### Ocular surface epithelial AHR mediates TCDD toxicity

Because AHR functions in a highly cell type- and tissue-specific manner (Anderson et al., 2022, Nebert et al., 2000, Omiecinski et al., 1990), we wanted to first determine if it was responsive to TCDD in the embryonic eyelids. For this objective, we assessed the expression of the prominent AHR target gene *Cyp1a1*, encoding Cytochrome P450 1a1. Most abundant CYP1A1 expression was found in the suprabasal epithelia on the exterior side of the eyelids in E15.5 embryos collected from pregnant dams treated with TCDD (50 µg/kg body weight) on E11.5. The expression was also detectable, albeit less abundantly, in the inferior epithelia and the stroma of exposed embryonic eyelids (Fig. 1A). As expected, CYP1A1 expression was undetectable in the eyelids of vehicle-treated embryos.

**Figure 1.**
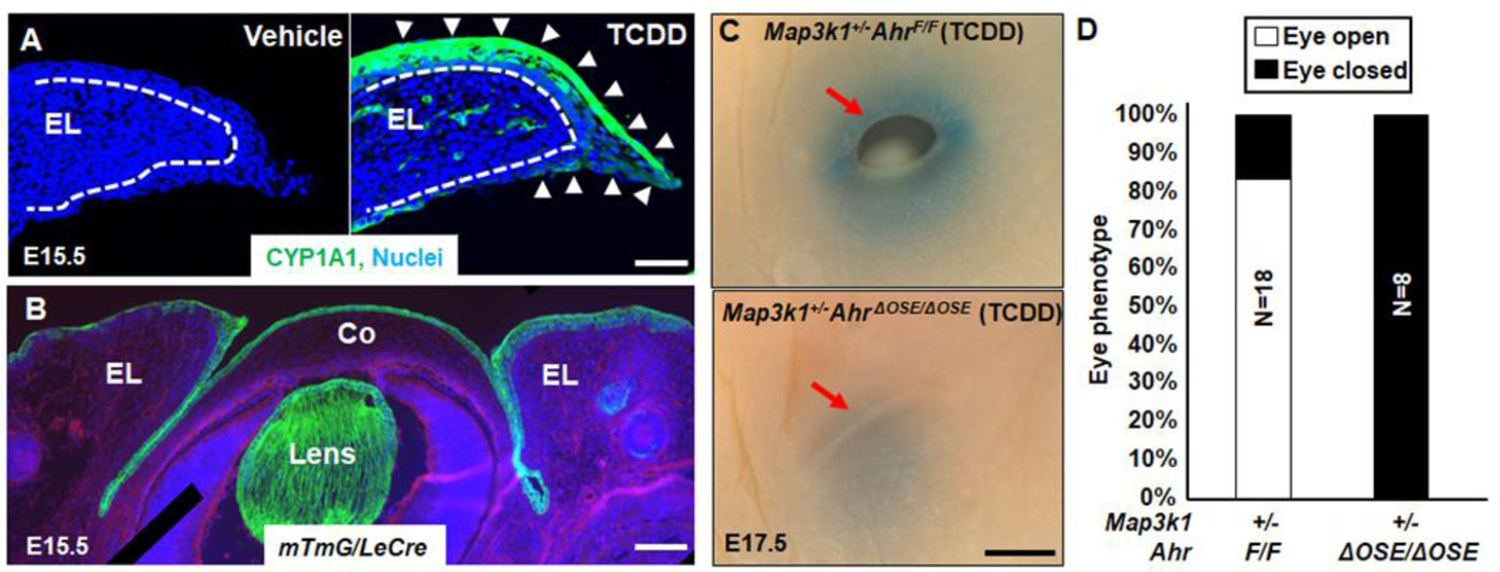
Eyelid epithelial AHR is required for TCDD toxicity in embryonic eyelid closure. (**A**) Immunofluorescent staining using anti-CYP1A1 (green) and Hoechst (blue) for nucleus of the eyelids in wild type E15.5 embryos with or without TCDD exposure (50 ug/kg body weight) at 11.5 days. Dash lines mark basement membrane of the eyelid epithelium; white arrows point at CYP1A1 positive staining. (**B**) Hoechst (blue) staining for nucleus of eyes of the E15.5 *mTmG*/*Le-Cre* embryos. Cre expressing, i.e., GFP positive, cells were located at the ocular surface epithelium (OSE) of the eyelid (EL), cornea (Co) and Lens. Pups exposed at E11.5 to TCDD (250 µg/kg body weight) were collected at E17.5, and their eyes (**C**) were photographed, red arrow, eyelid leading edge, and (**D**) quantification of eyes displaying open eye phenotype in *Map3k1^+/-^Ahr^F/F^*(N=18) and *Map3k1^+/-^Ahr^ΔOSE/ΔOSE^* (N=8) fetuses. N= number of pups of the indicated genotype. Scale bars, 50 µm in A, 100 µm in B, and 500 µm in C.

To address the role of the epithelial AHR in TCDD toxicity, we knocked out the *Ahr* gene in the developing ocular surface epithelium (OSE) by crossing *Ahr^F/F^* with *Le-Cre* mice. The *Le-Cre* transgene drives Cre recombinase expression in the developing OSE (shery-Padan et al., 2000). When tested in the *mTmG* reporter embryos, *Le-Cre* mediated GFP expression specifically in the epithelium, but not the stroma of the eyelid, cornea, and lens (Fig. 1B). The *Ahr^F/F^Le-Cre*, i.e., OSE-specific *Ahr* knockout (*Ahr^ΔOSE^*), were subsequently used to make *Map3k1^+/-^Ahr^F/F^* and *Map3k1^+/-^Ahr^ΔOSE^* compound mutants, both of which had normal eyelid closure in the absence of TCDD exposure. When dams were treated with high-dose (250 µg/kg) TCDD, 15 out of 18 (84%) *Map3k1^+/-^Ahr^F/F^* E17.5 fetuses exhibited the open-eye phenotype, whereas none of *Map3k1^+/-^Ahr^ΔOSE/ΔOSE^* fetuses (N=8) had the phenotype (Figs. 1C and 1D).

Another piece of evidence suggesting AHR mediated TCDD toxicity was the observations that all *Map3k1^+/-^* pups on a congenic C57BL/6 (B6) background developed the eyelid closure defects when exposed to 50 µg/kg TCDD, but the *Map3k1^+/-^Ahr^F/F^* pups showed the defects only when treated with 200 µg/kg or higher doses (Fig. S1). The differences in susceptibility to TCDD are likely the result of strain specific *Ahr* gene polymorphisms that give rise to AHR proteins with different ligand binding affinities (Walisser et al., 2004). The *Ahr* gene in B6 strain carries *Ahr^b1^* alleles that encode a high-affinity receptor, whereas the *Ahr* locus in *Ahr^F^* mice is derived from 129/SvJ strain that carries *Ahr^d^* alleles encoding a low-affinity receptor (Ema et al., 1994). Thus, the ligand binding affinity of AHR determines sensitivity of the eyelids to TCDD toxicity.

### Activation of the TCDD-AHR axis induces EGFR signaling

As the TCDD-AHR axis exerts biological effects through transcriptional regulation of gene expression (Hoffer et al., 1996), we set out to determine TCDD-AHR-dependent transcriptome in HaCaT, an aneuploid immortal human epithelial cell line (Boukamp et al., 1988). These cells possess a functional TCDD-AHR signaling, because TCDD induces a marked *CYP1A1* mRNA that is abolished by treatment with CH223191, an AHR antagonist (Fig. S2A). RNA-seq of the HaCaT cells treated with TCDD and CH22391 led to the identification of a total of 3,600 genes differentially expressed in a manner dependent on the TCDD-AHR pathway. Among these, 455 genes were up- or down-regulated by 2-fold or above (Figs. S2B and S2C). Ingenuity Pathway Analyses (IPA)^Tm^ showed that cell cycle regulation and xenobiotic AHR signaling were the top TCDD-AHR up-regulated functions, consistent with that reported previously (Sartor et al., 2009, Nebert et al., 2000) (Fig. S2D). On the other hand, epithelial-to-mesenchymal transition in development was identified as a top TCDD-AHR repressed function.

Further analyses of the transcriptome using the Upstream Regulator and Causal Network, which are designed to elucidate the biological causes and probable downstream effects (Kramer et al., 2014), led us to identify EGFR signaling as a target of the TCDD-AHR pathway. Specifically, TCDD induced the expression of EGFR upstream regulators, such as ERBB2, EGF and HRAS, and perturbed EGFR gene networks, including ERBB, tesevatinib, CUDC101 and lapatinib (Figs. 2A and 2B). The expression of several EGFR target genes, such as *IER3, EREG, EGR1, FOSL1, DUSP4* and *ETV5*, was found to be induced by TCDD and the induction was abolished by the AHR antagonist (Fig. 2C).

**Figure 2.**
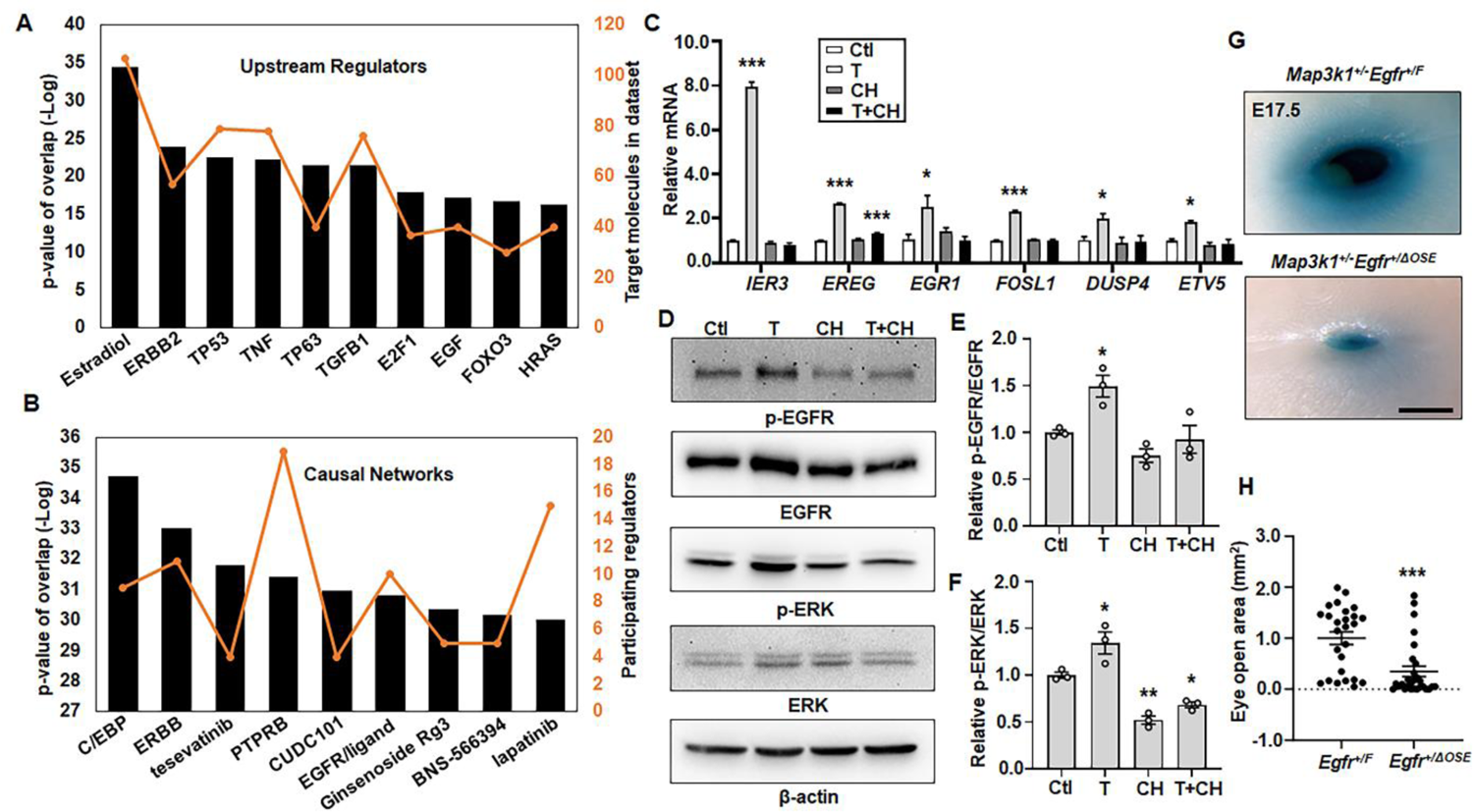
The TCDD-AHR axis activates EGFR signaling. The TCDD-AHR axis up-regulated genes were subjected to Ingenuity Pathway Analyses to identify (**A**) top upstream regulators and (**B**) top causal networks, with relative *p*-value (black bars) and number of participating molecules (orange line). (**C**) Selective gene expression was measured by qRT-PCR using RNA isolated from HaCaT cells treated with vehicle (DMSO, Ctl), TCDD (T, 10 nM), CH223191 (CH, 10 µM), and TCDD plus CH223191 (T+CH). Relative expression was calculated using the housekeeping gene *GAPDH* as an internal control and compared to that in Ctl set as 1. (**D**) Western blot analyses of the EGFR pathway activity using antibodies for phosphor (p) and total EGFR and ERK, and β-actin as a loading control. Quantification of (**E**) p-EGFR/EGFR and (**F**) p-ERK/ERK ratio of the immunoblotting data, with ratio in Ctl set as 1. The TCDD-exposed *Map3k1^+/-^Egfr^+/F^* (N=13) and *Map3k1^+/-^Egfr^+/ΔOSE^* (N=14) fetuses were collected at E17.5 and their eyes were (**G**) photographed and representative images were shown. Scale bar, 500 µm, and (**H**) the areas of open eye were measured. Results were shown as mean ± SEM of at least three independent experiments (N≥3). \**p*<0.05, \*\**p*<0.01 and \*\*\**p*<0.001 were considered statistically significant compared to Ctl (in C, E and F) and TCDD-treated *Map3k1^+/-^Egfr^+/F^*group (in H).

To verify the effect of TCDD on EGFR signaling, we measured EGFR tyrosine 1173 phosphorylation, a hallmark of receptor activation (Thompson and Gill, 1985). TCDD-treated HaCaT cells had a slight (1.5-fold) yet persistent increase of p-EGFR in comparison to vehicle-treated cells, and the increase was abolished by the AHR antagonist (Figs. 2D and 2E). Similarly, the phosphorylation of ERK, a downstream activation target of EGFR (Martinelli et al., 2017), was induced by TCDD in a manner dependent on AHR (Figs. 2D and 2F).

We also tested genetically whether the ocular surface EGFR contributed to the eyelid toxicity of TCDD using the *Map3k1^+/-^Egfr^+/F^*and *Map3k1^+/-^Egfr^+/ΔOSE^* pups, which had normal eyelid closure in the absence of TCDD exposure. After exposed to TCDD *in utero*, most of the EGFR wild type (*Egfr^+/F^*) *Map3k1^+/-^* fetuses developed the open-eye defects; however, the *Egfr^+/ΔOSE^Map3k1^+/-^* fetuses, in which EGFR expression in OSE was expected to be reduced by 50%, were less defective (Fig. 2G). The severity of the open-eye defect, i.e., size of eye opening, varied among individuals, but was overall significantly milder and decreased by 64% in *Map3k1^+/-^Egfr^+/ΔOSE^* compared to that in *Map3k1^+/-^Egfr^+/F^* fetuses (Fig. 2H). Hence, TCDD acts at least partially through the epithelial EGFR to disrupt eyelid closure.

### EGFR-ERK and MAP3K1-JNK form an inhibitory loop

Treatment of HaCaT cells with AG1478, a specific EGFR inhibitor, abolished the basal and TCDD-induced phosphorylation of EGFR and ERK, but conversely, it markedly increased JNK phosphorylation (Fig. 3A and 3B). Similarly, treatment of cells with PD98059, an ERK inhibitor, diminished the p-ERK, but induced the p-JNK (Figs. 3C and 3D). While these observations suggested that the EGFR-ERK pathway led to the repression of JNK activity, we found that inhibition of JNK by SP600125 induced the p-ERK (Figs. 3C and 3E), suggesting a mutual inhibitory relationship of JNK and ERK.

**Figure 3.**
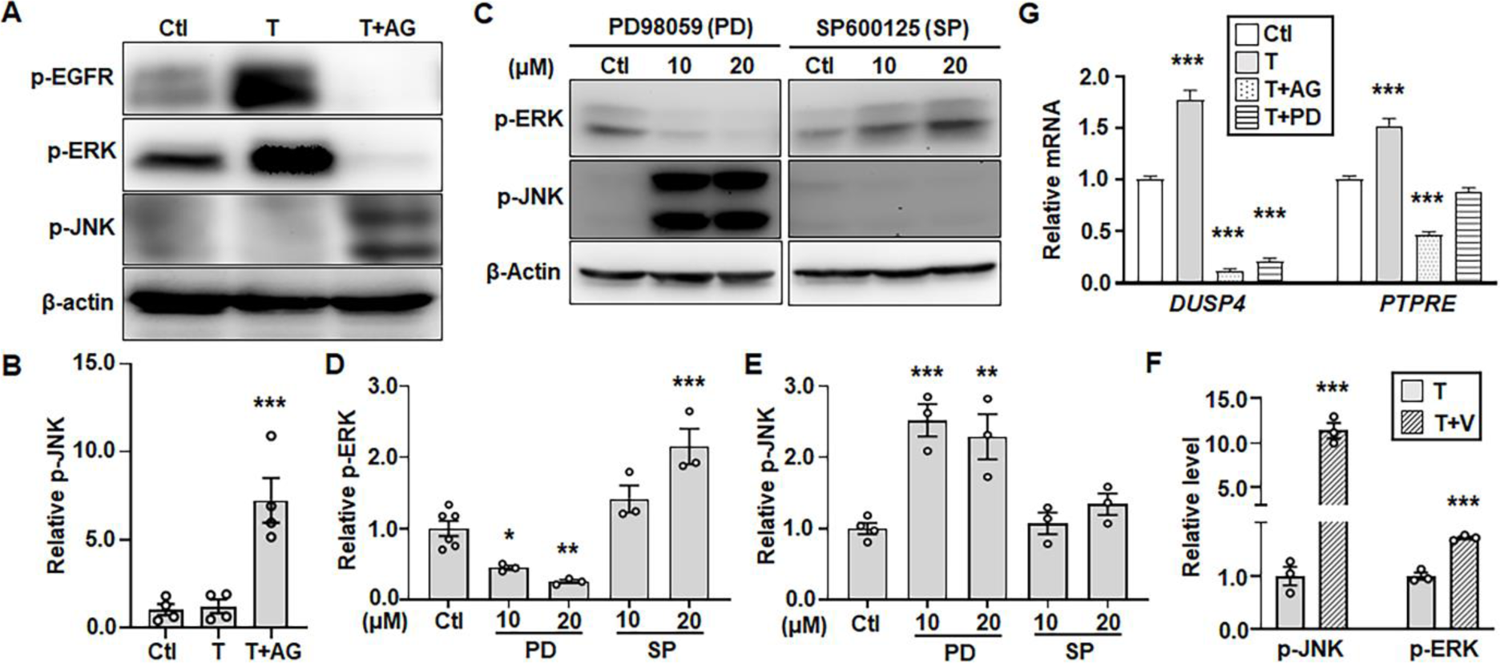
The EGFR-ERK pathway inhibits JNK activities. HaCaT cells treated with vehicle (DMSO, Ctl) and TCDD (T, 10 nM) in the presence or absence of the EGFR inhibitor, AG1478 (AG, 10 µM), were (**A**) subjected to immunoblotting for p-EGFR, p-ERK, p-JNK and β-actin, and (**B**) quantification for p-JNK using β-actin as a loading control. HaCaT cells treated with different concentrations of an ERK inhibitor, PD98059 (PD), and a JNK inhibitor, SP600125 (SP) were (**C**) examined by immunoblotting for p-ERK, p-JNK and β-actin, and values of (**D**) p-ERK and (**E**) p-JNK were quantified and compared to that of β-actin. (**F**) p-JNK, p-ERK and β-actin were examined by immunoblotting in lysates of HaCaT cells treated with TCDD (T, 10 nM) in the presence and absence of phosphatase inhibitor Na_3_VO_4_ (V, 0.5 mM). Values of relative p-JNK and p-ERK were quantified using β-actin. (**G**) Expression of *DUSP4* and *PTPRE* was examined by qRT-PCR in HaCaT cells treated with vehicle (DMSO, Ctl) and TCDD (T, 10 nM) in the presence and absence of 10 µM AG and PD. Relative mRNA of *DUSP4* and *PTPRE* was calculated using *GAPDH* as an internal control. Value of Ctl sample was set as 1. Data were shown as mean ± SEM of at least three independent experiments (N≥3). \**p*<0.05, \*\**p*<0.01 and \*\*\**p*<0.001 were considered significantly different compared to Ctl (in B, D, E and G) and TCDD-treated cells (in F).

When the HaCaT cells were treated with a non-specific phosphatase inhibitor Na_3_VO_4_ (V) (Aguilar et al., 2009, Chen and Tan, 1998), there was a significant increase of p-JNK by 12-fold and p-ERK by 2-fold (Fig. 3F). Based on these observations, we asked whether TCDD induced protein phosphatases to inhibit JNK signaling (Ip and Davis, 1998). Searching the transcriptomics, we found that the TCDD-AHR pathway induced the expression of several phosphatase genes, including *DUSP4*, *ALPPL2*, *ALPP* and *PTPRE* (Fig. S3A and Fig. S3B). Among them, the dual specificity phosphatase 4 (DUSP4) and protein tyrosine phosphatase epsilon (PTPRE), are potent enzymes in dephosphorylation and inactivation of JNK (Akimoto et al., 2009, Xu et al., 2023). Their expression was induced by TCDD in a manner also dependent on EGFR and ERK (Fig. 3G), suggesting that TCDD may act via EGFR-ERK to induce phosphatases for JNK repression.

As MAP3K1 is upstream of JNK signaling (Xia et al., 2000), we next assessed the involvement of MAP3K1 in the EGFR/ERK-JNK crosstalk. For an *in vitro* assessment, we made stable HaCaT cells in which MAP3K1 expression was either down-regulated with a small hairpin RNA, i.e., MAP3K1-deficient (shRNA), or specifically up-regulated with the sgRNA/CRISPR/dCas9 Synergistic Activation Mediator (SAM) system, i.e., MAP3K1-competent (SAM) (Fig. S3C). In these cells, we showed that increased MAP3K1 expression in the SAM cells led to elevated p-JNK, whereas decreased MAP3K1 in shRNA cells was associated with higher levels of p-EGFR and p-ERK (Figs. 4A and 4B). For the *in vivo* assessment, we examined the eyelids of wild type and *Map3k1*-null E15.5 embryos. Compared to the wild type embryos, the *Map3k1^-/-^* embryos had reduced p-JNK but elevated p-ERK in the suprabasal epithelium near the eyelid leading edge (Fig. 4C). Quantification of the immunostaining signals revealed a 45% reduction of p-JNK but 36% increase of p-ERK in *Map3k1^-/-^* versus wild type eyelids (Figs. 4D and 4E). Together, the *in vitro* and *in vivo* data suggest a feed-forward inhibitory loop, in which EGFR-ERK represses MAP3K1-JNK, and the repression of MAP3K1-JNK potentiates EGFR and ERK activities that may in turn further inhibit JNK.

**Figure 4.**
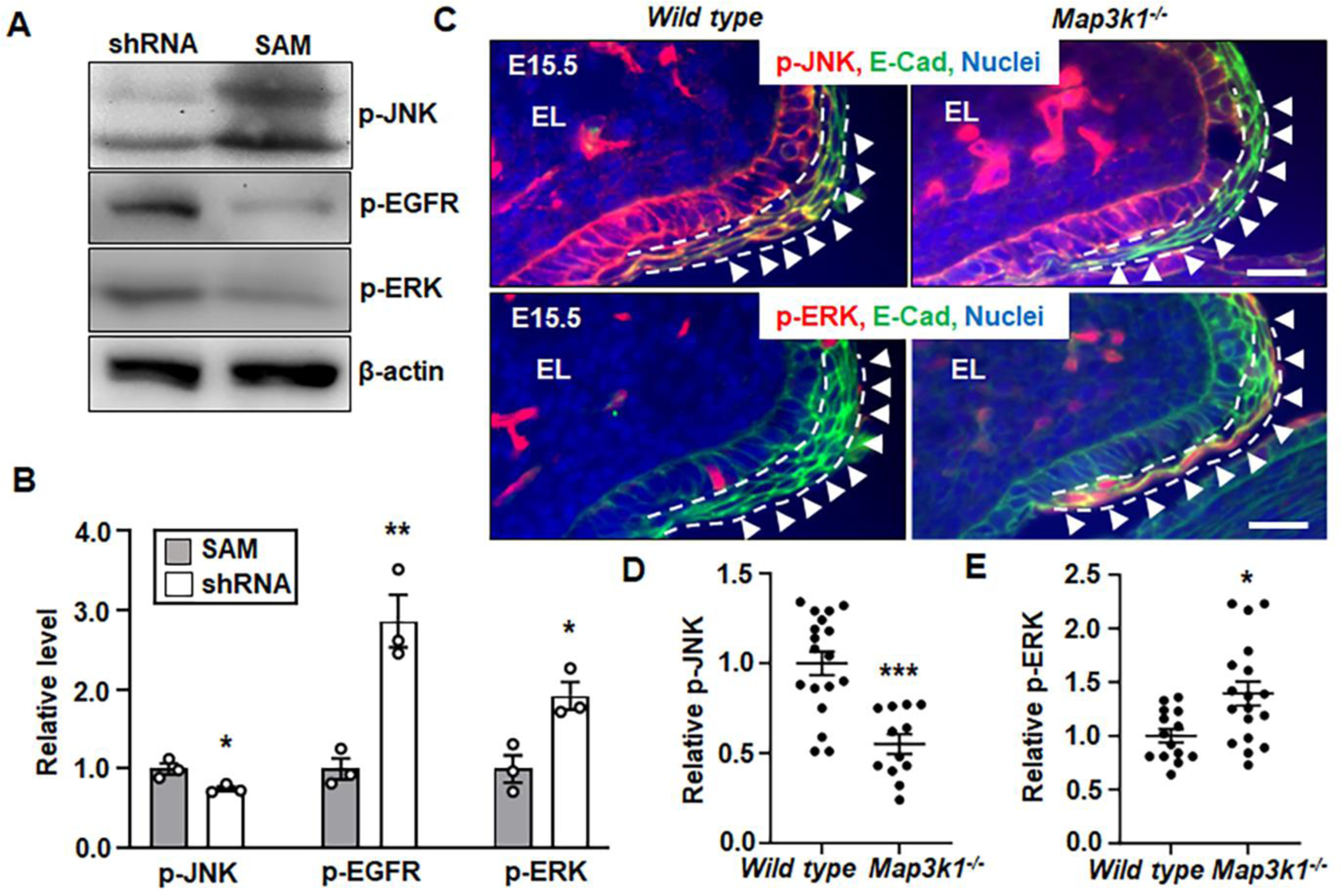
The MAP3K1-JNK and EGFR-ERK pathways inhibit each other. Lysates of the shRNA HaCaT and SAM HaCaT cells were examined by Western blotting for (**A**) p-JNK, p-EGFR, p-ERK, and β-actin, and (**B**) quantification of p-JNK, p-EGFR, and p-ERK using β-actin as a loading control. Levels in SAM HaCaT were set as 1. Data represented three independent experiments (N=3) and were shown as mean ± SEM. (**C**) Immunofluorescence staining of wild type and *Map3k1^-/-^* E15.5 embryonic eyelids with anti-p-JNK (red, top panels) and anti-p-ERK (red, bottom panels), co-stained with anti-E-cadherin (green) that marks epithelial membrane, and Hoechst (blue) labels nuclei. Representative Images were shown, scale bars, 50 µm. The (**D**) p-JNK, and (E) p-ERK, in the suprabasal epithelial cells, marked with dash lines, of the eyelid leading edge (arrowheads) were quantified and compared to that in wild type, set as 1. At least three sections (N≥3) per embryo and three embryos (N≥3) of each genotype from different litters were examined. Data were shown as mean ± SEM. \**p*<0.05, \*\**p*<0.01 and \*\*\**p*<0.001 were considered significantly different from SAM cells (in B) and wild type embryos (in D and E).

The feed-forward signaling loop might enable a substantial amplification of the small effects elicited by either TCDD or *Map3k1* heterozygosity when the two conditions were combined. In support of this possibility, we found that while the TCDD treatment (TCDD-treated wild type) or the *Map3k1* heterozygosity (vehicle-treated *Map3k1^+/-^*) did not significantly change p-JNK and p-ERK from those in vehicle-treated wild type, the TCDD-exposed *Map3k1^+/-^*eyelids had a marked increase of p-ERK and a significant decrease of p-JNK (Figs. 5A-5D).

**Figure 5.**
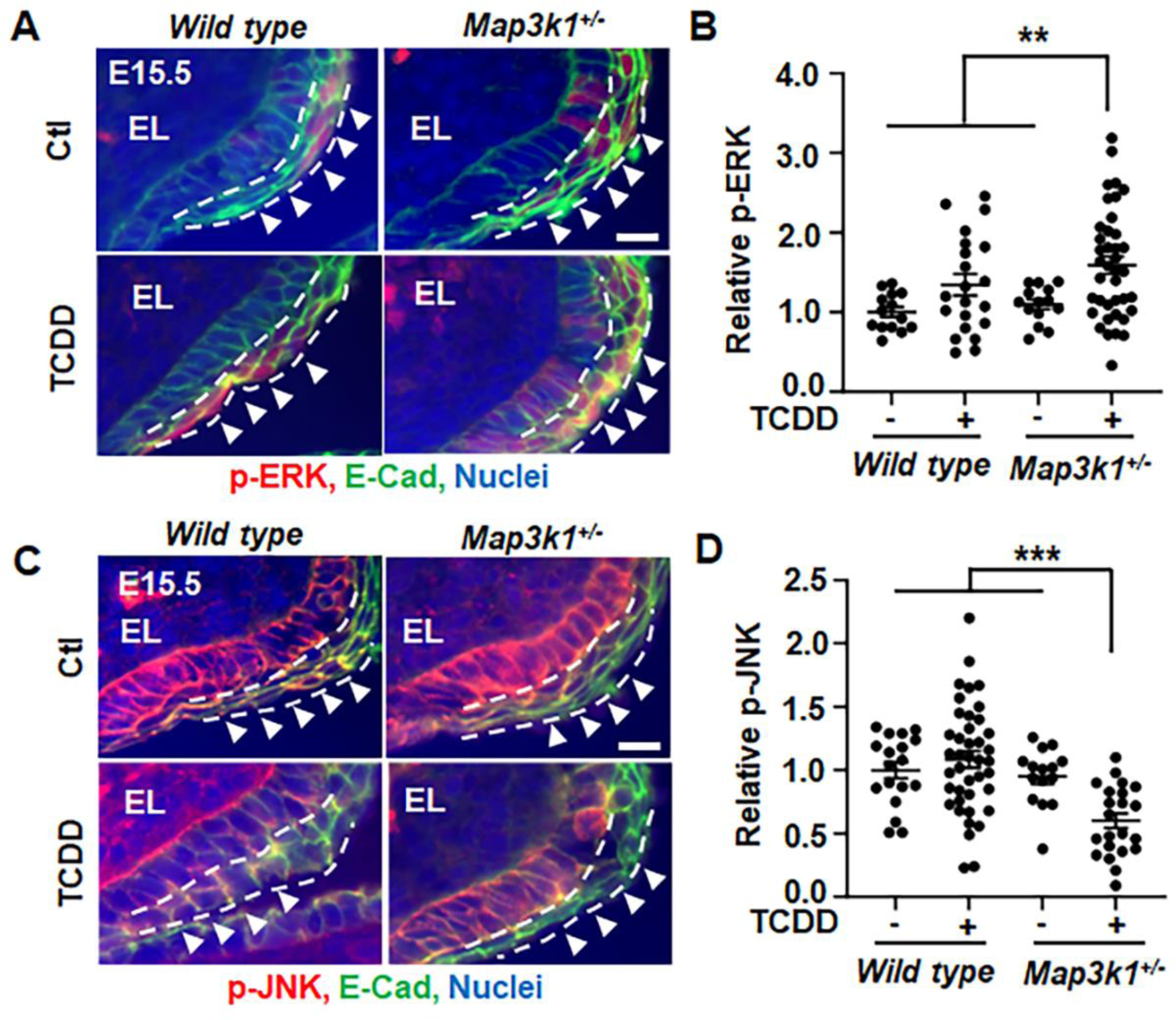
TCDD plus *Map3k1* heterozygosity tilt the balance of JNK and ERK. Eyelid tissues of wild type and *Map3k1^+/-^* E15.5 embryos with or without TCDD (50 µg/kg body weight) exposure were subjected to immunohistochemistry for (**A** and **B**) p-ERK (red) and (**C** and **D**) p-JNK (red). E-cadherin (green) and Hoechst (blue) were markers of epithelial membrane and nuclei, respectively. Representative images captured with a fluorescent microscopy were shown. The (B) p-ERK and (D) p-JNK in the suprabasal epithelium of the eyelid leading edge (arrowheads), marked with dash lines, were quantified. At least three sections (N≥3) per embryo and three embryos (N≥3) per genotype/treatment conditions were examined. Results were shown as mean ± SEM. \*\**p*<0.01, \*\*\**p*<0.001 represents significantly different from the unexposed groups and TCDD-treated wild type group (in B, D). Scale bar, 50 µm (A and C).

### JNK deficiency is the molecular basis of GxE interactions in tissue closure defects

The significant decrease of p-JNK in TCDD-exposed *Map3k1^+/-^* eyelids and the fact that both the *Map3k1^+/-^Jnk1^-/-^* and the *Jnk1* and *Jnk2* double gene knockout mice display the EOB phenotype (Weston et al., 2004, Takatori et al., 2008) led us to postulate that JNK deficient is responsible for the eyelid closure defects. To test this proposition, we assessed statistically the relationships between JNK activity and eyelid closure. To this effect, we measured the eyelid epithelial-specific p-JNK in embryos with normal eyelid development, under the wild type, *Map3k1^+/-^* and TCDD-exposed wild type conditions; we also measured p-JNK in embryos associated with the open-eye defects, under *Map3k1^-/-^*and TCDD-exposed *Map3k1^+/-^* conditions. The collective p-JNK data were subjected to receiver operating characteristic (ROC) analyses in a logistic regression model. The analyses revealed an inverse correlation between the levels of p-JNK and the possibility of the eyelid closure defects (Fig. 6A). Specifically, the probability of developing the defects was low when p-JNK was above an arbitrary threshold, but the probability was high (>55%) when p-JNK was below the threshold. The levels of p-JNK in embryos under TCDD combined with *Map3k1* heterozygosity fell largely below the threshold.

**Figure 6.**
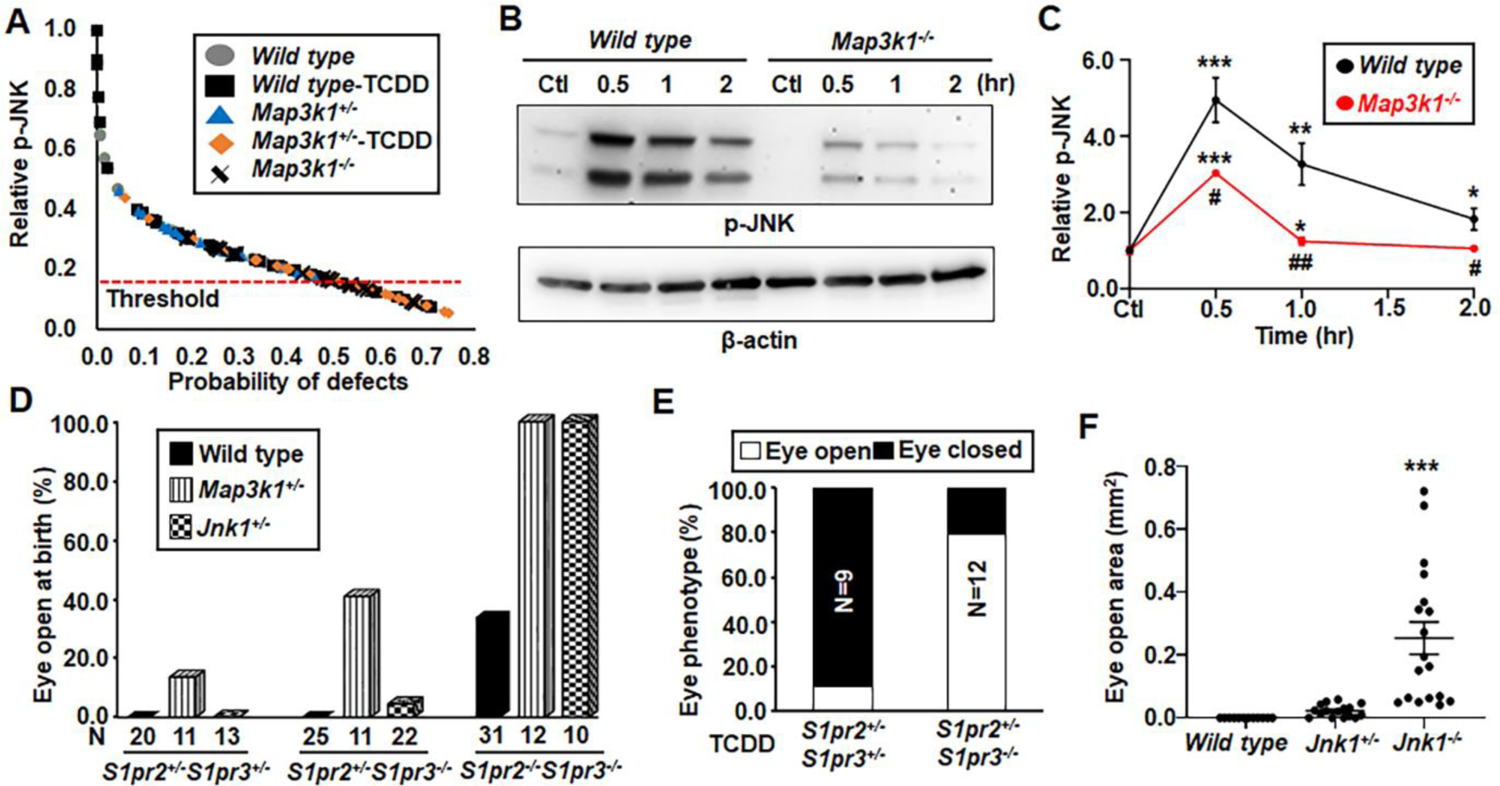
Crosstalking of TCDD and the S1PR-MAP3K1-JNK1 pathway for embryonic eyelid closure. (**A**) The levels of p-JNK in the eyelid epithelium were collected from E15.5 embryos of different genetic and exposure conditions as indicated. Data were analyzed by ImageJ software, and the relationship between the relative p-JNK level and the probability of the open-eye defects was calculated using a logistic regression model with ROC analysis. Probability of defect was significant higher when p-JNK was below 0.17. (**B**) The wild type and *Map3k1^-/-^* keratinocytes treated with vehicle (DMSO, Ctl) or 20 μM S1P for different time as indicated and p-JNK and β-actin were examined with Western blotting. (**C**) Quantification of p-JNK using β-actin as a loading control. Data are mean ± SEM of at least three independent experiments (N≥3). The *S1pr2*- and *S1pr3*-compound mutant pups under *Wild type, Map3k1^+/-^* and *Jnk1^+/-^*genetic backgrounds as indicated (**D**) without and (**E**) with *in utero* exposure to 50 ug/kg TCDD on E11.5 were collected at E17.5. The pups were examined for the eyelid open/close. N= number of pups of the indicated genotype. (**F**) The open eye areas were measured in E17.5 wild type, *Jnk1^+/-^* and *Jnk1^-/-^* fetuses exposed to TCDD (50 µg/kg body weight) at E11.5. At least 8 pups (N≥8) under each condition as indicated were analyzed. Values in Ctl were set as 1. \**p*<0.05, \*\**p*<0.01, \*\*\**p*<0.001 were significantly different from Ctl of the same genotype (in C) and TCDD exposed wild type (in F). #*p*<0.05, ##*p*<0.01 was significantly different between genotypes under the same treatment condition (in C).

The S1P is a phospholipid signaling molecule that exerts biological effects by binding to and activating specific G protein coupled S1P receptors (S1PRs) on plasma membrane (Sternweis et al., 2007, Aarthi et al., 2011). Because the S1PR2 and 3 display redundant roles in the regulation of embryonic eyelid closure (Herr et al., 2013), we explored whether the S1P signal was upstream of the MAP3K1-JNK pathway. We treated the wild type and *Map3k1^-/-^* keratinocytes with S1P for 30 – 120 min and examined JNK phosphorylation. Compared to the robust induction of p-JNK by S1P in the wild type cells, the induction was significantly less, reduced by more than 50% in the *Map3k1^-/-^* cells (Figs. 6B and 6C), suggesting that the majority of S1P signals for JNK activation were mediated through MAP3K1.

Genetic crossing, in which genes belong to the same molecular pathways display non-complementation for the EOB phenotype, is a powerful tool to map the signaling pathways *in vivo* (Takatori et al., 2008, Yook et al., 2001). Using this approach, we investigated the relationships between S1PR, MAP3K1 and JNK1. Consistent with observations of others (Herr et al., 2013), we found that a portion (30%) of *S1pr2/3* double knockouts developed the EOB phenotypes, but neither the *S1pr2^+/-^/S1pr3^+/-^* double heterozygous nor the *S1pr2^+/-^/S1pr3*^-/-^ triple allele deletion had the defects (Fig. 6D). The *S1pr*-mutant offspring, however, had higher incidence of the EOB phenotypes when crossed with either *Map3k1*- or *Jnk1*-knockout mice. Under the *Map3k1*^+/-^ genetic conditions, 13% of *S1pr2^+/-^/S1pr3^+/-^*, 41% in *S1pr2^+/-^/S1pr3*^-/-^ and 100% in *S1pr2/S1pr3*-double knockouts displayed the EOB defects. The *Jnk1*^+/-^ background had also increased the incidence of the EOB defects to 6% in *S1pr2^+/-^/S1pr3*^-/-^ and 100% in *S1pr2/3* double knockout pups. The genetic data strongly suggest the existence of a S1PR-MAP3K1-JNK pathway in embryonic eyelid closure.

If the endpoint of this pathway, i.e., JNK activity, is responsible for eyelid closure, we expect that genes contributing to JNK activity can modify TCDD toxicity, as seen in the *Map3k1^+/-^* pups. To test this prediction, we exposed pregnant dams carrying *S1pr2/3*-mutant embryos to TCDD and examined the open eye phenotypes in prenatal pups. TCDD induced the open-eye defects in 11% of *S1pr2^+/-^/S1pr3^+/-^* pups and 80% of *S1pr2^+/-^/S1pr3*^-/-^ pups (Fig. 6E). In comparison, all un-exposed pups of the same genotype had closed eyelids. We also tested this prediction in *Jnk1* gene knockout mice. In the absence of TCDD exposure, the *Jnk1^+/-^* and *Jnk1^-/-^* mice were born with closed eyelids (Kuan et al., 1999); however, after *in utero* exposure to TCDD, they displayed an open-eye defect (Fig. 6F). The severity of the defect inversely correlated with the number of *Jnk1* allele presented in the exposed fetuses - the wild type pups had normal eyelid closure, the *Jnk1* heterozygous pups had small eye openings, but the *Jnk1^-/-^* pups had apparent and large eye open defects. The results indicate that JNK dose-dependently antagonizes TCDD toxicity in eyelid development.

### The GxE interactions lead to aberrant epithelial morphogenesis and differentiation

To understand the biological consequences of the GxE interactions, we compared gene expression signatures in MAP3K1-competent (SAM HaCaT) cells and TCDD-treated MAP3K1-deficient (shRNA HaCaT) cells. There were approximately 24% genes differentially expressed by more than 2-fold in the comparison. Among these genes, 748 (24%) were up-regulated in TCDD-treated shRNA cells, whereas 2310 genes (76%) were down-regulated by TCDD combined with MAP3K1 deficiency (Fig. S4A). When further filtered based on AHR dependency, 82 genes (11%) were induced, and 79 (3%) genes were repressed by TCDD combined with MAP3K1 deficiency. Enrichment analyses of these genes identified Keratinocyte differentiation as one of the top functions up regulated and Cell morphogenesis as the top function downregulated by TCDD plus MAP3K1 deficiency (Fig. S4B and S4C).

The pathway analyses showed that either the TCDD-AHR pathway or the MAP3K1 deficiency repressed cell morphogenesis, but TCDD combined with MAP3K1 deficiency further repressed this function (Fig. S5A). Similarly, the morphogenetic gene expression was further decreased when the genetic and environmental factors combined (Fig. S5B). Some of morphogenetic genes, such as *FGFR2* and *FZD3,* have been implicated in embryonic eyelid closure (Huang et al., 2009, Fuhrmann, 2008), while some others, such as *RAC3* and *NEK3*, *DKK1*, *NOG*, *SMO*, *DAB2IP, PTPRZ1* and *GPM6A*, seem to belong to the RHO, WNT, TGFβ/BMP, SHH and MAPK morphogenetic pathways (Geh et al., 2011, Gage et al., 2008, Huang et al., 2009, Kuracha et al., 2013, Weston et al., 2003). Many other repressed genes, such as *CDH8, EFNA5/B3, EPHA4, ANOS1, TUBA1A, FLRT2 AMIGO1, POF1B, STMN1, SEMA6B* and *WDR19*, have a role in cell adhesion, cytoskeleton reorganization and migration.

A hallmark of epithelial morphogenesis is the establishment of cell polarity, where the Par3-Par6-αPKC complex and the acetyl-tubulin accumulate at distinct location to facilitate cell shape change, elongation and migration (Gibson and Perrimon, 2003, Chen and Zhang, 2013). In the embryonic eyelids, we found that the polarity markers, i.e., PAR6 and acetyl-tubulin, were readily detectable in the suprabasal epithelia leading to the eyelid tip protrusion in wild type and *Map3k1^+/-^* E15.5 embryos. However, these markers were absent in TCDD-exposed *Map3k1^+/-^* embryos (Figs. 7A and 7B).

**Figure 7.**
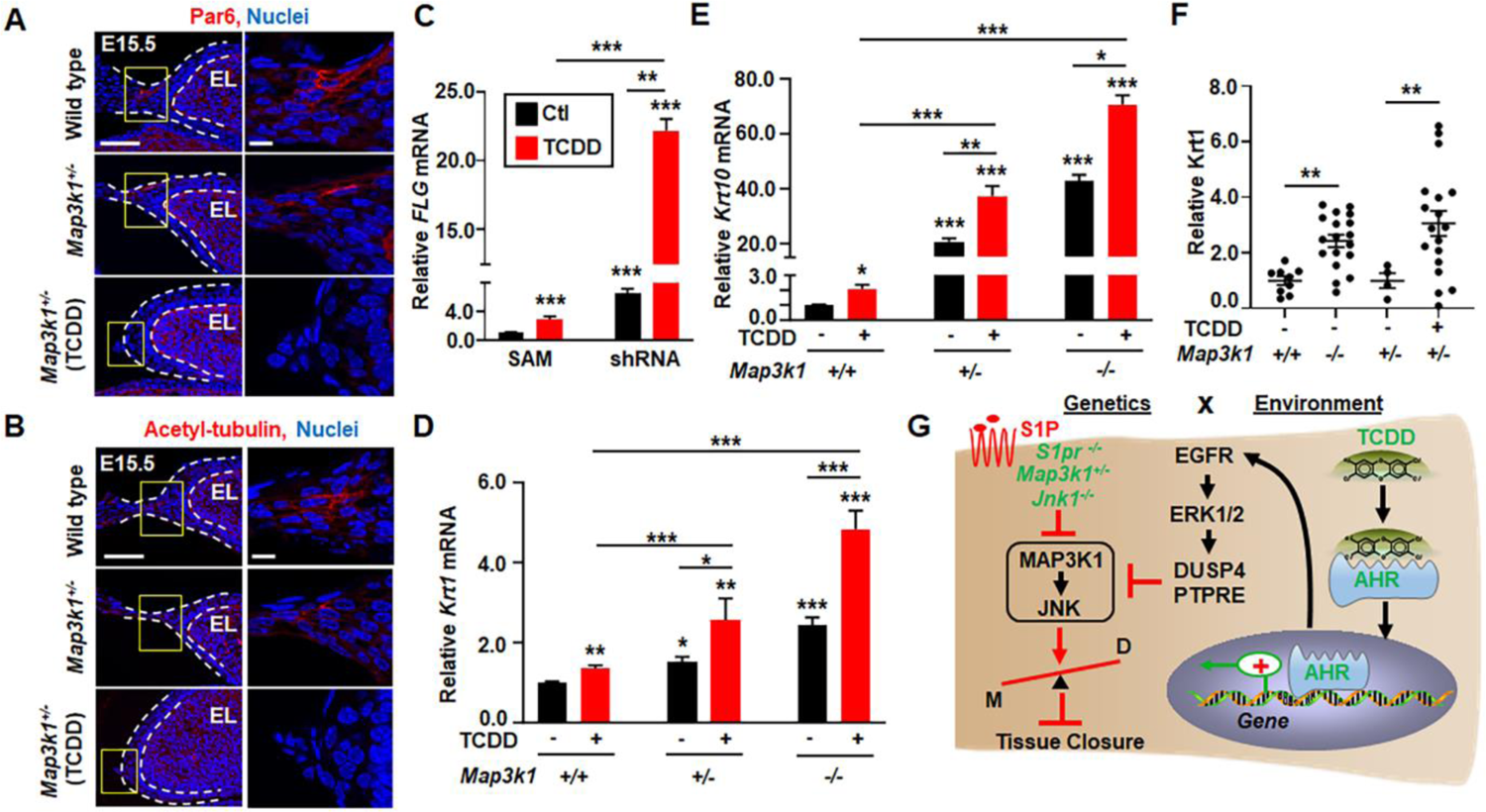
TCDD and *Map3k1* loss-of-function disrupt cell polarity and potentiate epithelial terminal differentiation. Eyelids of E15.5 wild type, *Map3k1^+/-^*, and TCDD treated *Map3k1^+/-^*embryos were subjected to immunohistochemistry using (**A**) anti-Par6 and (**B**) anti-acetylated-tubulin, detecting apical epithelial polarity and Hoechst labeling nuclei. Dash lines mark the eyelid epithelium. (**C**) The expression of *Filaggrin* (*FLG*), a marker of terminally differentiated keratinocytes in the granular and cornified epidermis, examined by qRT-PCR in shRNA HaCaT and SAM HaCaT treated with 10 nM TCDD or vehicle (DMSO, Ctl). The relative *FLG* mRNA was calculated using the housekeeping gene *GAPDH* as an internal control. *FLG* levels in SAM (Ctl) were set as 1, Expression of (**D**) *Krt1* and (**E**) *Krt10*, markers of the differentiating suprabasal keratinocytes, were examined in wild type, *Map3k1^+/-^*, and *Map3k1^-/-^* keratinocytes treated with 10 nM TCDD or vehicle (DMSO). Relative expression was calculated using the housekeeping gene *GAPDH* as an internal control and compared to levels in wild type keratinocytes set as 1. (**F**) The E15.5 embryos of wild type, *Map3k1^-/-^*, and *Map3k1^+/-^* with or without TCDD exposure were examined by immunohistochemistry for Krt1. Staining signals were quantified, and relative expression was calculated. Data are mean ± SEM of at least three independent experiments (N≥3) or at least three eye sections (N≥3) per embryo and three embryos (N≥3) from different litters. \**p*<0.05, \*\**p*<0.01, \*\*\**p*<0.001 were considered statistically significant compared to wild type and SAM untreated or as indicated. (**G**) Graphic illustration of the proposed GxE interactions model in eyelid closure defects. The environmental factor TCDD activates the AHR to induce gene expression and activate the EGFR-ERK pathways. EGFR signaling in turn induces phosphatases to inhibit the S1PR-MAP3K1-JNK pathway. This effect of TCDD is trivial and insufficient to induce the eyelid phenotype. However, in the presence of gene mutations, i.e., *S1pr2/3^-/-^, Map3k1^+/-^* and *Jnk1^-/-^*, which also slightly attenuate MAP3K1-JNK signaling, the effect of TCDD is largely amplified. As the results, the GxE interactions significantly inhibit JNK, accelerate differentiation (D) and impede morphogenesis (M) of the epithelium, leading to defective tissue closure.

The gene expression analyses also revealed that TCDD plus MAP3K1 deficiency (shRNA-TCDD vs SAM) accelerated keratinocyte differentiation (Fig. S5C and S5D). Many of the differentiation genes were up-regulated by TCDD-AHR or MAP3K1 deficiency, and their expression was further potentiated when the two conditions combined (Fig. S5D). Some of the genes, such as *S100A7* (Kizawa et al., 2011), *SPRR1A* (Heikinheimo et al., 2015) and *FLG* (Hoober and Eggink, 2022), are well-known markers of terminal differentiation of the epithelium.

To validate the GxE effects on differentiation, we examined the expression of *FLG,* encoding a component of the epidermal differentiation complex, in the genetically modified HaCaT cells. The basal *FLG* expression in MAP3K1-deficient (shRNA) cells was 6.9-fold of that in MAP3K1-competent (SAM) cells, whereas the expression was further increased to 23.4-fold above the basal levels in shRNA cell treated with TCDD (Fig. 7C). The epithelial differentiation was also examined in wild type, *Map3k1^+/-^* and *Map3k1^-/-^* cultured mouse keratinocytes. These cells maintain basal keratinocyte characteristics but spontaneously undergo terminal differentiation to the spinous epithelial cells (Wang et al., 2021b). The expression of *Keratin 1* (*Krt1*) and *Krt10*, products of which dimerize to form keratin intermediate filaments in the differentiated spinous epidermis, was relatively low in wild type cells, but significantly increased in *Map3k1^+/-^*, and further elevated in *Map3k1^-/-^* cells (Figs. 7D and 7E). TCDD treatment also potentiated the expression of these differentiation markers. Notably, the levels of *Krt1* and *Krt10* expression in TCDD-treated *Map3k1^+/-^*cells reached to the levels like that in *Map3k1^-/-^*cells. Similar findings were made in the embryonic eyelids. The expression of KRT1 protein was very low in the eyelid epithelium of wild type and *Map3k1^+/-^* E15.5 embryos; however, the expression was increased by 2.4-fold in that of *Map3k1^-/-^* embryos and 3.1-fold in TCDD-treated *Map3k1^+/-^*embryos (Fig. 7F). Taken together, our results show that TCDD and MAP3K1 deficiency, individually and in combination, accelerate epithelial terminal differentiation and impede cell polarity in the embryonic eyelids.

## Discussion

Here we show that TCDD activates the ocular surface epithelial AHR to impair embryonic eyelid closure in the *Map3k1^+/-^* mice. Tracking the molecular footprint of TCDD, we identify TCDD-AHR-regulated gene expression, leading to a moderate yet persistent activation of the EGFR-ERK pathway and the induction of protein phosphatases, which may in turn repress JNK signaling. The repression of JNK by TCDD is mild and insufficient to block eyelid closure; however, it becomes substantial in the presence of genetic perturbations. Specifically, *Map3k1* heterozygosity combined with TCDD further reduce p-JNK and cause the open-eye defects (Fig. 7G). We have also shown that S1P/S1PR is upstream of the MAP3K1-JNK pathway and that attenuation of the pathway activity by gene mutations, such as *S1pr2/3* and *Jnk1* knockout, also potentiates TCDD toxicity in eyelid morphogenesis. These results indicate that the crosstalk of genetic and environmental risk factors is mediated through an interactive signaling network, a mechanism likely extrapolatable to many diseases resulting from the GxE interactions in.

The GxE effects attribute primarily to the crosstalk between the MAP3K1/JNK and EGFR/ERK pathways. Under certain experimental settings, ERK and JNK are known to mutually inhibit each other to maintain a balanced signaling output, which is crucial for cell fate decisions (Junttila et al., 2008, Bluthgen and Legewie, 2008, Dong and Bode, 2003). We find a similar inhibitory relationship that extends to MAP3K1 and EGFR, upstream of JNK and ERK, respectively. As such, activation of EGFR/ERK by TCDD leads to the repression of JNK activity, whereas MAP3K1 knockdown/knockout also activates EGFR/ERK and represses JNK. As the result, the trivial effects on JNK repression and ERK activation by either the TCDD or the *Map3k1* heterozygosity become significantly amplified when the two conditions are combined. Our data, together with the findings that the *Jnk1/Jnk2* double knockout mice and the Ras-transgenic overexpression mice also exhibit the EOB defects (Weston et al., 2003, Burgess et al., 2010), suggest adverse imbalance of ERK/JNK signaling leads to impairment of eyelid closure. Alleviation of the imbalance through genetic perturbations, such as *Ahr ^ΔOSE^* and *Egfr^ΔOSE^*, that attenuate ERK signaling, reduces TCDD-induced eyelid defects. Conversely, aggravation of the imbalance through genetic ablation of *S1pr2/3* and *Jnk1* that attenuate JNK signaling potentiates the effects of TCDD.

JNK signaling is an evolutionarily conserved mechanism for developmental tissue closure. While it is required for dorsal closure and thorax closure through coordinating cell shape change in *Drosophila* (Jacinto et al., 2000, Martin-Blanco et al., 2000), it regulates gastrulation through controlling tissue elongation in *Xenopus* (Carron et al., 2005, Kim and Han, 2005). In mice, compound knocking out the functionally redundant *Jnk1* and *Jnk2* results in defective closure of the neural tube, the optic fissure, and the eyelid (Weston et al., 2003). In this work, we have additionally shown that JNK dose-dependently contributes to tissue closure. There is an inverse correlation between the number of *Jnk1* allele or the amount of p-JNK with the severity of eyelid closure defects. In this context, JNK may phosphorylate paxillin to regulate intercellular junctions interconnected with polarity (Huang et al., 2003, You et al., 2013). It is reasonable to speculate other genetic and environmental insults that target JNK signaling could potentially increase the risks of developing tissue closure abnormalities.

TCDD is known to activate the EGFR-ERK pathway and accelerate epithelial differentiation (Hansen et al., 1997, Sibilia and Wagner, 1995, Sutter et al., 2011, Kennedy et al., 2013, Sutter et al., 2020, Sutter et al., 2023, Fernandez-Salguero et al., 1996, Puga et al., 2005, Choi et al., 2006, Patel et al., 2006), an effect that may lead to promotion of precocious eyelid opening in postpartum pups (Theobald and Peterson, 1997, Gray et al., 1997, Miettinen et al., 2004, Dominey et al., 1993) and development of chloracne, a hyperkeratotic skin disorder, found in humans exposed to high dose TCDD (Sutter et al., 2011, Muenyi et al., 2014, Loertscher et al., 2001). This effect of TCDD, however, is insufficient to block embryonic lid closure. We show that like TCDD, *Map3k1* deficiency also induces EGFR/ERK activation and epidermal differentiation. In the embryonic eyelid epithelium and cultured keratinocytes, TCDD combined with *Map3k1* deficiency have an additive or perhaps a synergistic effect on potentiating differentiation. Similar observations have been made during *in vitro* stem cell-to-epithelium differentiation, where TCDD plus *Map3k1* deficiency further accelerate basal to spinous differentiation (Wang et al., 2022). The premature differentiation may reduce flexibility of the eyelid tip epithelial cells, thereby preventing them to undergo shape change for lid closure (Byrne et al., 1994). Hence, the small effects on cell activities by each agent are exacerbated by the GxE interactions for a detrimental outcome.

Defective embryonic eyelid closure leads to developmental abnormalities of the ocular adnexa with phenotypes comparable to human congenital disorders, such as eyelid ptosis and strabismus (Meng et al., 2014). The etiology for these diseases is complex and not fully understood (Lorenz, 2002, Oystreck et al., 2012, Finsterer, 2003). Our results suggest the existence of multifactorial etiology, where the genetics and the environment act as rheostats for each other to induce the defect, but each condition alone is less, or not at all, destructive. Based on these findings, it is reasonable to predict that monogenic, digenic, and polygenic variants that affect the AHR/EGFR/ERK-S1PR/MAP3K1/JNK network will increase or decrease the toxic effects of TCDD and other environmental AHR ligands. Because embryonic eyelid closure shares similar morphogenetic activities with other tissue closure events, such as neural tube closure, palate fusion and ventral body wall closure, the signaling mechanisms underlying the GxE interactions described here may have important implications for the causation of abnormalities in other anatomical tissue closures. Identifying the genetic susceptibility of chemical toxicity is a crucial first step to uncover risk factors and improve personalized protection.

## Materials and methods

### Mouse strains, genotyping, and chemical treatment

The wild type C57BL/6J strain and the global green fluorescence Cre reporter (ROSA^mTmG^) strain were purchased from the Jackson Laboratory; the *Map3k1*, *Jnk1* and *S1pr2/3* mutant strains were described before (Xia et al., 2000, Zhang et al., 2003, Herr et al., 2013). The *Egfr^F/F^, AhrF^/F^* and Le-Cre mice were gifts from Drs. Threadgill, Bradfield and Ashery-Padan, respectively (Maklad et al., 2009, Walisser et al., 2004, shery-Padan et al., 2000). All mice were backcrossed for > 12 generations to obtain congenic C57BL/6J background. Genomic DNA PCR was used to determine genotype using gene-specific primers synthesized by Integrated DNA Technologies (Coralville, IA).

*In utero* exposure was done by oral gavage treatment of the pregnant dams on gestation day 11.5 with either corn oil (vehicle) or 50 to 500 µg/kg TCDD dissolved in corn oil, as described before (Mongan et al., 2015). Embryos/fetuses were harvested at E15.5 or E17.5 for histology, immunohistochemistry, and phenotype evaluation. Mice were housed in the Laboratory Animal Medical Service facility at the University of Cincinnati, College of Medicine. Experimental procedures were approved by the Institutional Animal Care and Use Committee (Protocol 23-11-01-01). All procedures with mice handling were in adherence to Guide for the Care and Use of Laboratory Animals and NIH or MRC guidelines for animal welfare.

### Cells, culture reagents and chemical treatment

Mouse keratinocyte lines derived from wild type and *Map3k1^-/-^*mice were cultured and maintained in Keratinocyte Serum Free Medium without Calcium chloride (KSFM-Ca) as described before (Wang et al., 2021b). HaCaT, a spontaneously immortalized human keratinocyte line, was from ATCC and cultured in Dulbecco’s Modified Eagle’s Medium (DMEM, Gibco) supplemented with 10 % fetal bovine serum, 2 mM glutamine, 1% nonessential amino acids, penicillin (100 U⁄ml) and streptomycin (100 µg⁄ml) in 5% CO_2_ at 37°C incubator.

The MAP3K1 deficient (shRNA) HaCaT cells were generated by transduction of HaCaT cells with MAP3K1 shRNA lentiviruses. To make lentivirus, the lentiviral vectors purchased from Sigma (Clone ID, TRCN000000616) were co-transfected with packaging psPAX2 (Addgene #12260) and envelope pMD2.G (Addgene #12259) plasmids into 293T packaging cells (ATCC). The lentiviruses were used to infect HaCaT cells, and the transduced cells were selected with puromycin (3 mg/ml) to obtain stable HaCaT shRNA cells. MAP3K1-competent HaCaT cells were generated using CRISPR/Cas9 Synergistic Activation Mediator (SAM) system (Konermann et al., 2015). Plasmids for the SAM system (i.e., lenti-sgRNA (MS2)_puro, dCas9VP64, and MS2-P65-HSF1) were purchased from Addgene (Addgene, cat. #73797, 61425, and 61426). The single guide RNAs (sgRNAs) for *MAP3K1* were designed based on publicly available filtering tools (https://zlab.bio/guide-design-resources) (Table S1) and cloned into lenti-sgRNA (MS2) puro. HaCaT cells were transduced with lentiviruses (i.e., sgRNA, MS2 and dCas9) and selected with puromycin (3 mg/ml), blasticidin (10 mg/ml) and hygromycin (10 mg/ml) to generate stable HaCaT SAM cells.

Cells were treated with 10 nM TCDD in the presence or absence of 10 µM CH-223191 for 24 h or with 10 µM AG1478, 10-20 µM PD98059 and SP600125 for 8 h or with 20 µM S1P for different times and harvested for RNA and protein analyses. Cell culture media, reagents, chemicals, and inhibitors are listed in Table S2.

### Phenotype evaluation and immunohistochemistry

E17.5 fetuses were harvested, and the eyes were photographed with a Leica MZ-16FA dissecting microscope. The size of the eye open was determined by tracing the edge of the eyelids and measuring areas with ImageJ software. Pixel values were transferred to actual areas (mm^2^) based on the scale information of each image. The value is zero if the eyelids are fully closed.

For immunohistochemistry, E15.5 embryos were harvested and fixed with 4% paraformaldehyde at 4°C overnight, followed by embedding in O.C.T compound and freeze. Eye sections were subjected to immunohistochemistry with specific primary antibodies and fluorescent labelled secondary antibodies. Primary and secondary antibodies are listed in Table S3. Images were captured by the Zeiss Axioplan 2 fluorescence and confocal microscope equipped with AxioVision software. In E-cadherin-positive epithelial cell layers, the suprafacial layer was outlined, in which the p-JNK or p-ERK were measured. The mean intensity (total intensity/area) was determined with the ImageJ software; the relative p-JNK/p-ERK levels were calculated after background subtraction and compared to those in wild type samples, set as 1. For the threshold model, data of p-JNK and open-eye were analyzed by a logistic regression model with receiver operating characteristic (ROC) analyses.

### Western Blotting

Cells were lysed in RIPA buffer containing 20 mM Tris-HCl, pH 7.5, 150 mm NaCl, 1 mM EDTA, 1% Nonidet P-40, 0.5% sodium deoxycholate, 10 μg/ml aprotinin, 10 μg/ml leupeptin, 1 mM Na3VO4, 100 μM phenylmethylsulphonyl fluoride. Lysates (100 µg protein) were subjected to SDS-PAGE. Proteins were transferred to nitrocellulose membranes and incubated with antibodies. Antibodies for western blotting are listed in Table S3. Immunoblotting signals were captured with UVP imaging system. Data were quantified with ImageJ using densitometry normalized with β-actin as loading control. The relative protein level was compared to those in Ctl, TCDD treated or SAM cells, set as 1.

### RNA isolation, reverse transcription, and Real Time-PCR (RT-PCR)

Total RNA was isolated using PureLink RNA Mini Kit (12183025, Invitrogen). First-strand complementary DNAs (cDNAs) were synthesized by 0.5 μg of total RNA using SuperScript™ IV Reverse Transcriptase (Invitrogen) according to the manufacturer’s instruction. PCR reactions were carried out with Stratagene Mx3000P RT-PCR system (Agilent Technologies). Power SYBR™ Green PCR Master Mix (Applied Biosystems) was used as the detection format. The reaction was heated to 95°C for 10 min, followed by 40 cycles of denaturation at 95°C for 15 sec and annealing elongation at 60°C for 60 sec. Gene expression was calculated by comparative ΔΔCt-method normalized to a constitutively expressed housekeeping gene (*GADPH*). Data represent results of triplicates in at least three independent experiments. The sequences of PCR primers are listed in Table S4.

### RNA-seq

RNA quality was QC analyzed by Bioanalyzer (Agilent, Santa Clara, CA). RNA sequencing of biological triplicate samples was performed by the Genomics, Epigenomics and Sequencing Core at the University of Cincinnati using established protocols described previously (Wang et al., 2021b). Details were in the Supplementary Materials and Methods. Raw data and processed files were either uploaded to NCBI GEO database (GSE237130) and in the process of submission.

### Transcriptome analyses

To identify genes regulated by the TCDD-AHR pathways, we compare differential gene expressions in HaCaT cells treated for overnight with 1) vehicle, 2) 10 nM TCDD, 3) 10 µM AHR inhibitor CH223191, and 4) TCDD plus CH223191. Gene expression in samples 2) were compared with combined samples in 1), 3) and 4) to identify significantly differential expressed genes dependent on TCDD-AHR (3600 genes) (padj<0.05). The TCDD-AHR dependent genes were submitted to QIAGEN IPA for Pathway analysis, Upstream Regulators and Causal Networks. Heat maps were generated by using Heatmapper (Babicki et al., 2016).

To identify genes affected by MAP3K1 loss-of-functions, we compared gene expressions in shRNA vs SAM cells. The upregulated (log2 fold change >1, padj <0.05) and downregulated (log2 fold change <-1, padj <0.05) genes were subjected to functional enrichment analyses with the Metascape software (v3.5.20230501) (Zhou et al., 2019). To analyze the gene expression signatures of the gene-environment interactions, we compared expression in TCDD-treated shRNA HaCaT with un-treated SAM HaCaT; the data were further filtered for TCDD-AHR dependent genes. The resultant differential expressed genes were subjected to functional enrichment analysis using Metascape.

### Data quantification and statistical analysis

Quantification of the eye open phenotype, immunohistochemistry positive signals and immunoblotting band intensity were done by Image J software (ver. Fiji-2.1.1). All data are shown as mean ± standard error of the mean (SEM) based on at least three independent experiments and analyzed using Student’s two-tailed t-test or One-way analysis of variance following Dunnett’s test for multiple comparisons. #*p*, \**p*<0.05; ##*p*, \*\**p*<0.01; and \*\*\**p*<0.001 are considered statistically significant.

## Acknowledgements

The authors wish to thank Drs. David Threadgill at Texas A&M University, Christopher Bradfield at University of Wisconsin, Ruth Ashery Padan at Tel Aviv University, and Jerold Chun at Sanford Burnham Prebys Medical Discovery Institute for providing genetic modified mice, and Dr. Antonius Christianto for critical reading of the manuscript.

## Data availability

All relevant data can be found within the article and its supplementary information.

